# Identification of parallel and divergent optimization solutions for homologous metabolic enzymes

**DOI:** 10.1101/260554

**Authors:** Robert F. Standaert, Richard J. Giannone, Joshua K. Michener

## Abstract

Metabolic pathway assembly typically involves the expression of enzymes from multiple organisms in a single heterologous host. Ensuring that each enzyme functions effectively can be challenging, since many potential factors can disrupt proper pathway flux. These challenges are amplified when the enzymes are expressed at single copy from the chromosome. We have explored these issues using 4-hydroxybenzoate monooxygenase homologs heterologously expressed in *Escherichia coli.* Initial chromosomal enzyme expression was insufficient to support consistent growth with 4-hydroxybenzoate. Experimental evolution identified mutations that improved pathway activity. One set of mutations was common between homologs, while a second class of mutations was homolog-specific. Ultimately, we were able to identify a set of mutations that provided sufficient activity for growth with 4-hydroxybenzoate while maintaining or improving growth with protocatechuate. These findings demonstrate the potential for flexible, scalable chromosomal pathway engineering, as well as the value of directed evolution strategies to rapidly identify and overcome diverse factors limiting enzyme activity.

## 1.0 Introduction

Synthetic biologists frequently transfer metabolic pathways from native hosts into more-tractable production strains (Nielsen and Keasling, 2016). By combining enzymes from different organisms, they can construct novel pathways to produce valuable biochemicals (Galanie et al., 2015; Yim et al., 2011). However, combining diverse enzymes into complex pathways is challenging, since the interactions between enzymes within a pathway, or between a pathway and its host, can limit productivity (Kim and Copley, 2012; Michener et al., 2012). A common solution is to screen multiple enzyme homologs, with the goal of identifying a variant that lacks deleterious interactions (Bayer et al., 2009; Narcross et al., 2016). A deeper understanding of the reasons that enzyme homologs function poorly will increase the success rate of screening homolog libraries, and thereby allow faster and cheaper pathway assembly.

The catabolism of lignin-derived aromatic chemicals provides a representative example of this engineering challenge. The thermochemical depolymerization of lignin produces a complex mixture of aromatic compounds (Rodriguez et al., 2017). Though a process described as biological funneling, microbes can convert this low-value mixture of substrates into a small set of core intermediates and then into valuable products (Linger et al., 2014). However, no single microbe has yet been isolated that is capable of catabolizing and valorizing all the constituents of a typical mixture (Bugg et al., 2011). Instead, researchers seek to augment the catabolic capabilities of foundation organisms by introducing new metabolic pathways for conversion of recalcitrant substrates (Strachan et al., 2014). Changes in the feedstock composition and pretreatment strategy will alter the composition of the depolymerized mixture (Ragauskas et al., 2014), necessitating the construction and optimization of an entire suite of microbes, each specific to a particular substrate mixture. Consequently, the facile valorization of lignin-derived aromatics will require the ability to rapidly engineer metabolic networks using pathways from diverse microbes.

Heterologous metabolic pathways are frequently expressed from plasmids. These vectors are easy to construct, and the resulting high DNA copy numbers can compensate for low specific enzyme activity. However, plasmid-based pathways are unstable and difficult to scale for larger pathways, such as those that will be required for catabolism of a mixture of lignin-derived aromatic compounds (Jiang et al., 2013). Moving heterologous enzymes from plasmids to the chromosome increases stability and scaling, and due to new techniques is increasingly practical (Jiang et al., 2015). However, the reduced copy number increases the challenge of providing sufficient enzyme activity (Alonso-Gutierrez et al., 2017; Wang and Pfeifer, 2008).

In this work, we explored the challenges involved in extending a metabolic pathway using chromosomally-expressed enzyme homologs, focusing specifically on pathways for the catabolism of 4-hydroxybenzoate (4-HB) and protocatechuate (PCA) (Figure 1). These compounds are the core metabolites for catabolism of hydroxyphenyl (H) and guaiacyl (G) lignans, respectively, and therefore are important targets for engineering biological conversion of lignin components. We previously constructed and optimized a pathway in *E. coli* that converts PCA to succinate and acetyl-CoA, allowing growth with PCA as the sole source of carbon and energy (Clarkson et al., 2017). We then extended the pathway to convert 4-HB into PCA using a 4-hydroxybenzoate monooxygenase, *praI*, from *Paenibacillus* sp. JJ-1B (Kasai et al., 2009), allowing *E. coli* to grow with 4-HB as the carbon source, with limitations that we have now discovered. In the present work, we introduced an alternate 4-hydroxybenzoate monooxygenase homolog, *pobA*, from *Pseudomonas putida* KT2440 (Harwood and Parales, 1996; Jimenez et al., 2002). Using either of the monooxygenases, we selected mutant strains that grew efficiently with 4-HB as the sole source of carbon and energy, at the same rate as with PCA, and without the previous limitations.

**Figure 1:**
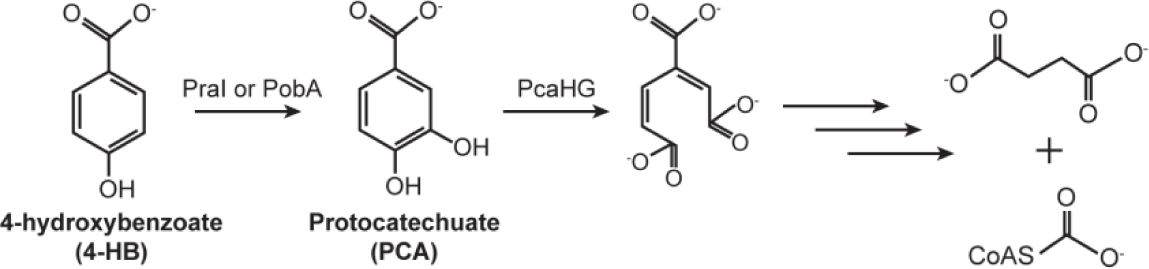
Catabolism of 4-HB. The 4-HB monooxygenases PraI and PobA convert 4-HB into PCA. The first step in PCA degradation is ring cleavage by the PCA 3,4-dioxygenase PcaHG, ultimately yielding succinate and acetyl-CoA.

Characterizing the resulting evolutionary solutions identified multiple interacting factors that limited 4-HB catabolism and revealed unexpected differences between enzyme homologs.

Similar RNA secondary structures, unintentionally introduced during codon optimization, initially decreased expression of both 4-HB monooxygenases. After resolving this issue, we found that the monooxygenases are not fully interchangeable, as different optimization solutions produced varied results with the two homologs. Ultimately, we identified pathway modifications for both homologs that allowed similar levels of growth with either PCA or 4-HB. These modifications included duplications of the core PCA degradation pathway and point mutations in the PCA/4-HB transporter, PcaK. Unexpectedly, these modifications had very different impacts on *pral* and *pobA* strains. Either allowed *pobA* strains to grow on 4-HB, but only the PcaK mutations allowed *pral* strains to grow. The results provide examples of the modifications required to optimize the function of chromosomally-expressed enzyme homologs, facilitating the rapid, predictable construction and debugging of novel metabolic pathways.

## 2.0 Materials and methods

### 2.1 Media and chemicals

All chemicals were purchased from Sigma-Aldrich (St. Louis, MO) or Fisher Scientific (Fairlawn, NJ) and were molecular grade. All oligonucleotides were ordered from IDT (Coralville, IA). *E. coli* strains were routinely cultivated at 37 °C in LB broth containing the necessary antibiotics (50 mg/L kanamycin or 50 mg/L spectinomycin). Growth assays with PCA and 4-HB were performed in M9 salts medium containing 300 mg/L thiamine and 1 mM isopropyl β-D-1-thiogalactopyranoside (IPTG). PCA and 4-HB were dissolved in water at 5 g/L, filter sterilized, and added at a final concentration of 1 g/L. The pH of the substrate stock solutions was not adjusted, as PCA oxidation in air occurred more rapidly at neutral pH.

### 2.2 Plasmid construction

Expression for sgRNA plasmids targeting chromosomal loci were constructed as described previously (Clarkson et al., 2017). Briefly, an inverse PCR was used to amplify the vector backbone. Overlapping oligonucleotides containing the new 20-nt targeting sequence were inserted by Gibson assembly and transformed into 10-β *E. coli* (NEB, Waltham, MA). Correct assembly was verified by Sanger sequencing.

### 2.3 Strain construction

Strain JME17 was previously optimized for growth with PCA. For growth with 4-HB, the appropriate 4-HB monooxygenase was introduced into the chromosome of JME17. Construction of JME50, expressing the 4-HB monooxygenase PraI, was described previously (Clarkson et al., 2017). Strain JME38 was constructed in a similar fashion, introducing a commercially-synthesized *pobA* expression cassette (Gen9, Cambridge, MA) into JME17. The promoter and terminator for *pobI* expression were chosen from previously characterized genetic parts (Chen et al., 2013; Kosuri et al., 2013). The expression cassette was integrated into the *gfcAB* locus using plasmid pJM168 (Jiang et al., 2015). To reconstruct evolved mutations, the mutant locus was amplified from the appropriate genomic DNA and introduced into the selected recipient strain in the same fashion. To introduce a second copy of *pcaHGBDC*, the expression cassette was amplified from JME17, combined with homology arms for *yiaU* by overlap-extension PCR, and transformed into JME17 together with plasmid pJM193. All modifications were verified by colony PCR and Sanger sequencing, or by whole-genome resequencing.

### 2.4 Experimental evolution

Parental strains were streaked to single colonies. Three colonies were grown to saturation in LB + 1 mM IPTG, then diluted 128-fold into M9 + 1 mM IPTG + 1 g/L 4-HB + 50 mg/L PCA and grown at 37 °C. When the cultures reached saturation, they were diluted 128-fold into fresh medium. Initially, the saturated cultures had a low optical density (~0.2). Over time, the final density increased. When the final density passed 0.8, the PCA concentration was reduced, first to 20 mg/L, then to 10 mg/L, 5 mg/L, and finally 0 mg/L. After a reduction in the PCA concentration, a culture often required one to two additional days to reach saturation. After 300 generations, each population was streaked to single colonies. Eight replicate colonies were picked for further characterization.

### 2.5 Growth rate measurements

Cultures were grown overnight to saturation in M9 + 1 mM IPTG + 2 g/L glucose. They were then diluted 100-fold into fresh M9 + IPTG containing the appropriate carbon source and grown as triplicate 100 μL cultures in a Bioscreen C plate reader (Oy Growth Curves Ab Ltd, Helsinki, Finland). Growth rates were calculated using CurveFitter software based on readings of optical density at 600 nm (Delaney et al., 2013).

### 2.6 Genome resequencing

Genomic DNA (gDNA) was prepared using a Blood and Tissue kit (Qiagen, Valencia, CA) according to the manufacturer’s directions. The gDNA was quantified using a Qubit fluorimeter (Thermo Fisher, Waltham, MA) and resequenced by the Joint Genome Institute on a MiSeq (Illumina, San Diego, CA) to approximately 100x coverage.

### 2.7 Proteomics

Parent, evolved and engineered *E. coli* strains were grown in triplicate and processed for LC-MS/MS analysis. Whole-cell lysates were prepared by bead beating in sodium deoxycholate lysis buffer (4% SDC, 100 mM ammonium bicarbonate, pH 7.8) using 0.15 mM zirconium oxide beads followed by centrifugation to clear debris. After measuring protein concentration via BCA (Pierce), samples were adjusted to 10 mM dithiothreitol and incubated at 95°C for 10 min to denature and reduce proteins. Cysteines were alkylated/blocked with 30 mM iodoacetamide during 20 min incubation at room temperature in the dark. Protein samples (250 ug) were then transferred to a 10 kDa MWCO spin filter (Vivaspin 500, Sartorius), cleaned up, and digested *in situ* with proteomics-grade trypsin as previously described (Clarkson et al., 2017). Tryptic peptides were then collected by centrifugation, SDC precipitated with formic acid, and precipitate removed from the peptide solution water-saturated ethyl acetate. Peptide samples were then concentrated via SpeedVac and measured by BCA.

Peptide samples were analyzed by automated 2D LC-MS/MS analysis using a Vanquish UHPLC plumbed directly in-line with a Q Exactive Plus mass spectrometer (Thermo Scientific) outfitted with a triphasic MudPIT back column (RP-SCX-RP) coupled to an in-house pulled nanospray emitter packed with 30 cm of 5 μm Kinetex C18 RP resin (Phenomenex). For each sample, 5 μg of peptides were loaded, desalted, separated and analyzed across two successive salt cuts of ammonium acetate (50 mM and 500 mM), each followed by 105 min organic gradient. Eluting peptides were measured and sequenced by data-dependent acquisition on the Q Exactive as previously described (Clarkson et al., 2017).

MS/MS spectra were searched against the *E. coli* K-12 proteome concatenated with relevant exogenous protein sequences and common protein contaminants using MyriMatch v.2.2 (Tabb et al., 2007). Peptide spectrum matches (PSM) were required to be fully tryptic with any number of missed cleavages; a static modification of 57.0214 Da on cysteine (carbamidomethylated) and a dynamic modification of 15.9949 Da on methionine (oxidized) residues. PSMs were filtered using IDPicker v.3.0 (Ma et al., 2009) with an experiment-wide false-discovery rate (assessed by PSM matches to decoy sequences) initially controlled at < 1% at the peptide-level. Peptide intensities were assessed by chromatographic area-under-the-curve using IDPicker’s embed spectra/label-free quantification option, and unique peptide intensities summed to estimate protein-level abundance. Protein abundance distributions were then normalized across samples and missing values imputed to simulate the MS instrument’s limit of detection. Significant differences in protein abundance were assessed by one-way ANOVA and pairwise T-test. Statistical significance was evaluated using a Benjamini-Hochberg false discovery rate of 5%.

### 2.8 qPCR

Gene copy numbers were determined by quantitative PCR using Phusion DNA Polymerase (NEB) and EvaGreen (Biotium, Fremont, CA) on a CFX96 Touch thermocycler (Bio-Rad, Hercules, CA). Whole cells from triplicate cultures were used as the DNA templates. Copy number was normalized to 16S rRNA gene copies.

### 2.8 Intracellular metabolite measurements

Strains were grown overnight in M9 + glucose. They were then diluted 100x into fresh M9 + glucose and regrown to mid-log phase. The cells were centrifuged and resuspended in fresh M9 + glucose containing the appropriate substrate at an optical density of ~3, 1 mL per replicate. Samples were incubated at 37 °C for 15 minutes, washed twice in M9 + glucose, transferred to a fresh tube, and stored at -80 °C until analysis.

Metabolites (4-HB and PCA) were analyzed by gas chromatography/mass spectrometry (GC/MS) as follows. Frozen cell pellets were thawed and resuspended in 100 μL of water by vortex mixing, and lysozyme was added to a final concentration of 100 μg/mL by addition of 5 μL of a 2 mg/mL stock solution. Cells were incubated at room temperature for 30 min and then acidified with 10 μL of 0.1 M HCl containing 10 μg/mL of 4-HB-d_4_ as an internal standard. Metabolites were extracted by adding 1.0 mL of ethyl acetate, vortex mixing for 2 x 10s, centrifuging (16,000 x g for 2 min), transferring 0.8 mL of the ethyl acetate layer to a 2-mL glass autosampler vial, and evaporating the solvent under a stream of argon. Metabolites were derivatized by adding 100 μL of *N, O*-bis(trimethylsilyl)trifluoroacetamide containing 1% chlorotrimethylsilane, sealing the vials, heating to 60 °C for 30 min, and diluting with 900 μL of *n*-hexane.

GC/MS analysis was performed using an Agilent 7890A gas chromatograph equipped with a 7693A automatic liquid sampler, an HP-5ms capillary column (30 m long × 0.25 mm inside diameter with a 0.25μm capillary film of 5% phenyl methylsilicone) and a 5975C mass-sensitive detector. Splitless injections of 1 μL were made at an inlet temperature of 270 °C with a 15-s dwell time (needle left in the inlet after injection), an initial column temperature of 50 °C and helium carrier gas at a constant flow of 1 mL/min. After 2 min at 50 °C, the temperature was ramped at 25 °C/min to 300 °C and held for 2 min. The detector was operated with a transfer line temperature of 300 °C, source temperature of 230 °C and quadrupole temperature of 150 °C. After a 5-min solvent delay, electron-impact mass spectra from 50-400 amu were collected continuously (~4/s) for the duration of the run. Relative amounts of metabolites were determined by integration of selected ion chromatograms for PCA(TMS)_3_, 4-HB(TMS)_2_ and 4-HB-d_4_(TMS)_2_ at m/z 193, 267 and 271, respectively, and then dividing by the area of the 271 peak (internal standard).

## 3.0 Results and Discussion

### 3.1 Pathway construction and characterization

We previously described the construction of *E. coli* strain JME50 (Clarkson et al., 2017), containing*pral* from *Paenibacillus* sp. JJ-1B (Kasai et al., 2009). To explore the different optimization challenges of enzyme homologs, we used a similar process in this work to construct JME38, introducing *pobA* from *Pseudomonas putida* KT2440 (Harwood and Parales, 1996; Jimenez et al., 2002). These two genes are 60% identical at the nucleotide level, and the associated enzymes have 54% amino acid identity.

We reported that JME50 could grow in minimal medium containing 4-HB as the sole source of carbon and energy, in contrast to the parental strain JME17 lacking*praI* (Clarkson et al., 2017). Upon further investigation, however, we discovered that this strain was unable to grow in minimal medium with 4-HB if it had previously been cultured in minimal medium with glucose or 4-HB as the carbon source (Supplementary Figure 1). The same was true of JME38: expression of a 4-hydroxybenzoate monooxygenase was necessary for growth with 4-HB, but was not sufficient for repeated propagation. Rather than screening more enzyme homologs, in the hopes of finding an enzyme with sufficient activity, we decided to understand why these enzymes failed to function effectively and to modify both the enzymes and host to enable efficient catabolism of these challenging carbon sources.

### 3.2 Experimental evolution identifies improved variants

To uncover the factors limiting growth with 4-HB, we serially passaged three replicate cultures each of strains JME38 and JME50 for 300 generations in minimal medium containing 1 g/L 4-HB. Initially, cultures were supplemented with 50 mg/L of PCA to allow slight growth of the parental strains. The PCA concentration was reduced over time and eliminated after generation 100. After 300 generations, we isolated eight mutant strains from each replicate culture and measured their ability to grow with PCA and 4-HB. We then selected one representative isolate from each replicate population for further characterization.

All six isolates grew at similar rates with 4-HB (Figure 2). Two of the isolates showed reduced growth with PCA, as compared to the parental strain. Changes in growth rate with glucose were minor. We then resequenced the genome of each isolate. All six isolates had mutations at the 5’ end of the *pral* or *pobA* coding sequence, five of which were silent mutations (Supplementary Table 4). Based on sequence coverage, all three *pobA* mutants were predicted to have duplications of the *pcaHGBDC* operon. Conversely, all three *pral* mutants had unique, non-synonymous mutations in the gene encoding the PCA/4-HB transporter, *pcaK.* Each isolate had one or two additional mutations in the genome that were unique to that isolate.

**Figure 2:**
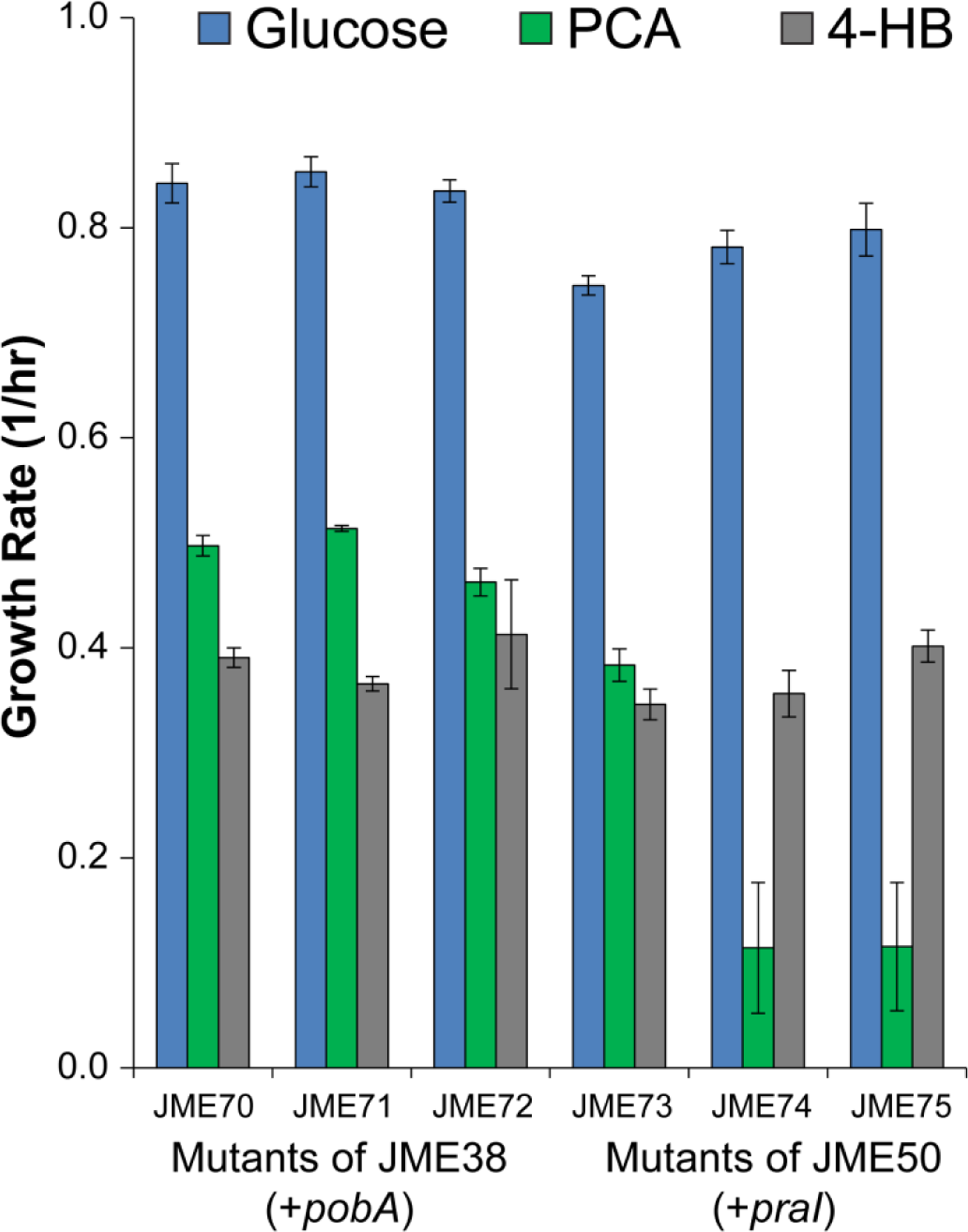
Evolved strains grow efficiently with 4-HB. Isolates from each of three replicate populations were assayed for growth in minimal medium with 2 g/L glucose, 1 g/L PCA, or 1 g/L 4-HB. Error bars show one standard deviation, calculated from three biological replicates.

For verification, we used quantitative PCR to measure the copy number of *pcaH* in the parental and evolved strains, since this is the first enzyme in the operon and PcaH and PcaG together catalyze the first step in PCA degradation (Figure 1B). The relative copy number of *pcaH* was two-to three-fold higher in the evolved isolates relative to the parental strain (Supplementary Figure 2A). To understand the effects of the*pcaHGBDC* duplication, we measured expression levels of the entire proteome in both the parental and evolved isolates (Supplementary Figure 2B). Consistent with our previous observations (Clarkson et al., 2017) and the qPCR results, we saw increased expression of PcaH and PcaG in the evolved strain (p<0.002).

### 3.3 Reconstruction identifies lineage-specific causal mutations

We introduced the silent *praI* and T388A *pcaK* mutations from JME73, individually and in combination, into JME50 to verify their effects. Neither mutation alone was sufficient for growth with 4-HB, while the double mutant was able to grow efficiently (Figure 3A). Similarly, we tested the silent *pobA* mutation from JME70 and an engineered duplication of the *pcaHGBDC* operon, alone or in combination. Only the double mutant was able to grow with 4-HB (Figure 3B). To understand why we saw different evolutionary outcomes in the two lineages, we then swapped the *pcaK* and *pcaHGBDC* mutations. The *pcaK* mutation supported rapid growth of a strain with *pobA*, while the *pcaHGBDC* duplication had minimal effect on a strain with *praI* (Figure 3C).

**Figure 3:**
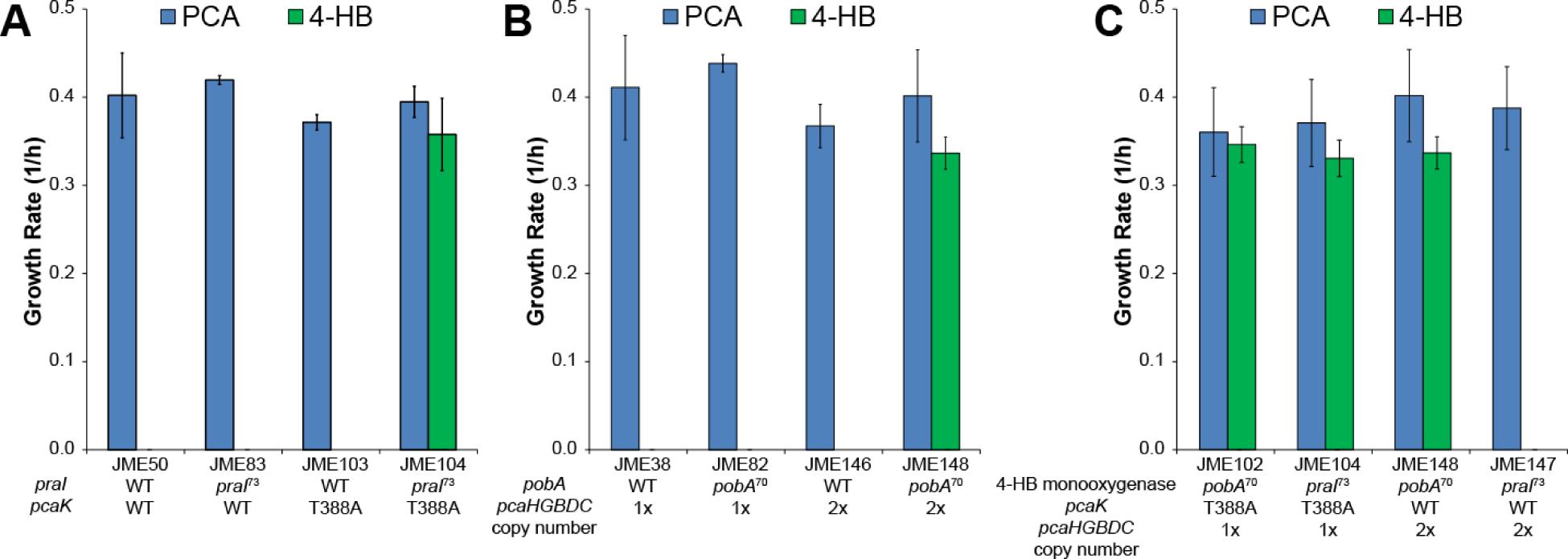
Two mutations are necessary for efficient growth with 4-HB. (A) Strains with either the wild-type or mutant alleles of *praI* and *pcaK* were grown with PCA or 4-HB as the sole source of carbon and energy. Mutation numbers for *praI* and *pobA* correspond to the strain from which those mutations were identified, as shown in Fig. 4. (B) As in A, except using strains with either the wild-type or mutant allele of *pobA* and one or two copies of *pcaHGBDC.* (C) As in A, except using strains with the mutant alleles of *praI* or *pobA* and either the mutant allele of *pcaK* or a second copy of *pcaHGBDC.* Error bars show one standard deviation, calculated from three biological replicates.

To understand the effect of the *pcaHGBDC* duplication, we measured the growth rate of strains JME17, with one copy of *pcaHGBDC*, and JME146, with a second copy, across a range of PCA concentrations (Supplementary Figure 3). While the growth rate of the two strains was similar at high PCA concentrations, JME17 grew more slowly at low PCA concentrations.

Insertion sequence-mediated duplications, such as those seen with *pcaHGBDC*, occur at much higher frequencies than specific point mutations (Andersson and Hughes, 2009). While strains with *pobA* can benefit from either mutations in *pcaK* or duplication of *pcaHGBDC*, we would expect the duplication to occur more often and therefore rise to fixation before a point mutation to *pcaK* would likely arise. Conversely, strains with *praI* were unable to benefit from the frequent duplications and instead waited for the rare mutations to *pcaK*.

### 3.4 Silent mutations increase expression of heterologous proteins

Mutations to *pobA* and *praI* that do not affect the primary structure of the protein were necessary for growth with 4-HB. To understand the effects of these mutations, we measured the expression of PobA and PraI in strains with wild-type and mutant alleles. In both cases, the mutations increased expression of the monooxygenase by more than three-fold (Figure 4, p<0.01).

**Figure 4:**
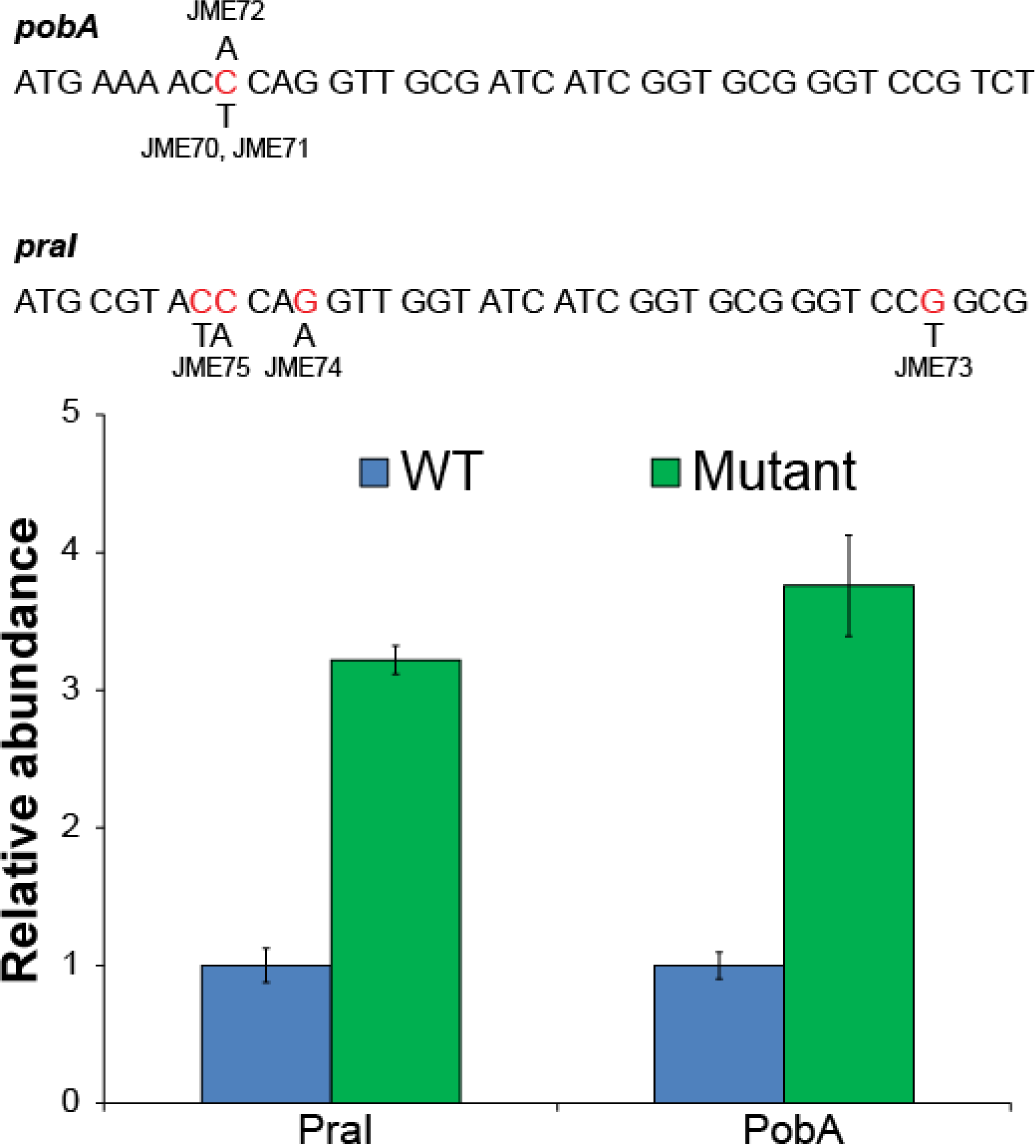
Silent mutations increase protein expression. All six evolved isolates had mutations to the 5’ end of the coding region of the monooxygenase gene, five of which were silent. The silent mutations present in JME70 and JME83 significantly increased abundance of the associated proteins, as measured by global proteomics. Error bars show one standard deviation, calculated from three biological replicates.

Secondary structures at the 5’ end of an mRNA can decrease protein abundance (Boël et al., 2016; Kudla et al., 2009). We calculated the predicted folding energy of the 5’ UTR and first 18 amino acids for each of the wild-type and mutant expression constructs. Each of the mutations increased the predicted local mRNA folding energy, i.e. destabilized the mRNA folding, by 1.4 to 4.1 kcal/mol. Changes in folding energy at this scale have been shown to be capable of producing changes in protein abundance of approximately three-fold (Kudla et al., 2009). The parallelism in mutations between *pobA* and *pral* likely results from a combination of high local amino acid identity combined with codon optimization. While the enzymes are 54% identical across their entire length, the first 18 amino acids are 78% identical. Codon optimization for *E. coli* eliminated any pre-existing nucleotide variation and yielded nucleotide sequences that were 83% identical across this region.

### 3.5 Mutations to*pcaK* affect transport of 4-HB

To understand how mutations to *pcaK* affect growth with 4-HB, we studied one particular mutation, T388A, in more detail. Expression of membrane proteins in a heterologous host can be difficult, and mutations to the protein may improve expression. Therefore, we measured changes in the expression of PcaK in otherwise-isogenic strains containing either the wild-type or mutant alleles of *pcaK.* Expression of PcaK was roughly two-fold higher for the mutant allele (Figure 5A, p<0.001).

**Figure 5:**
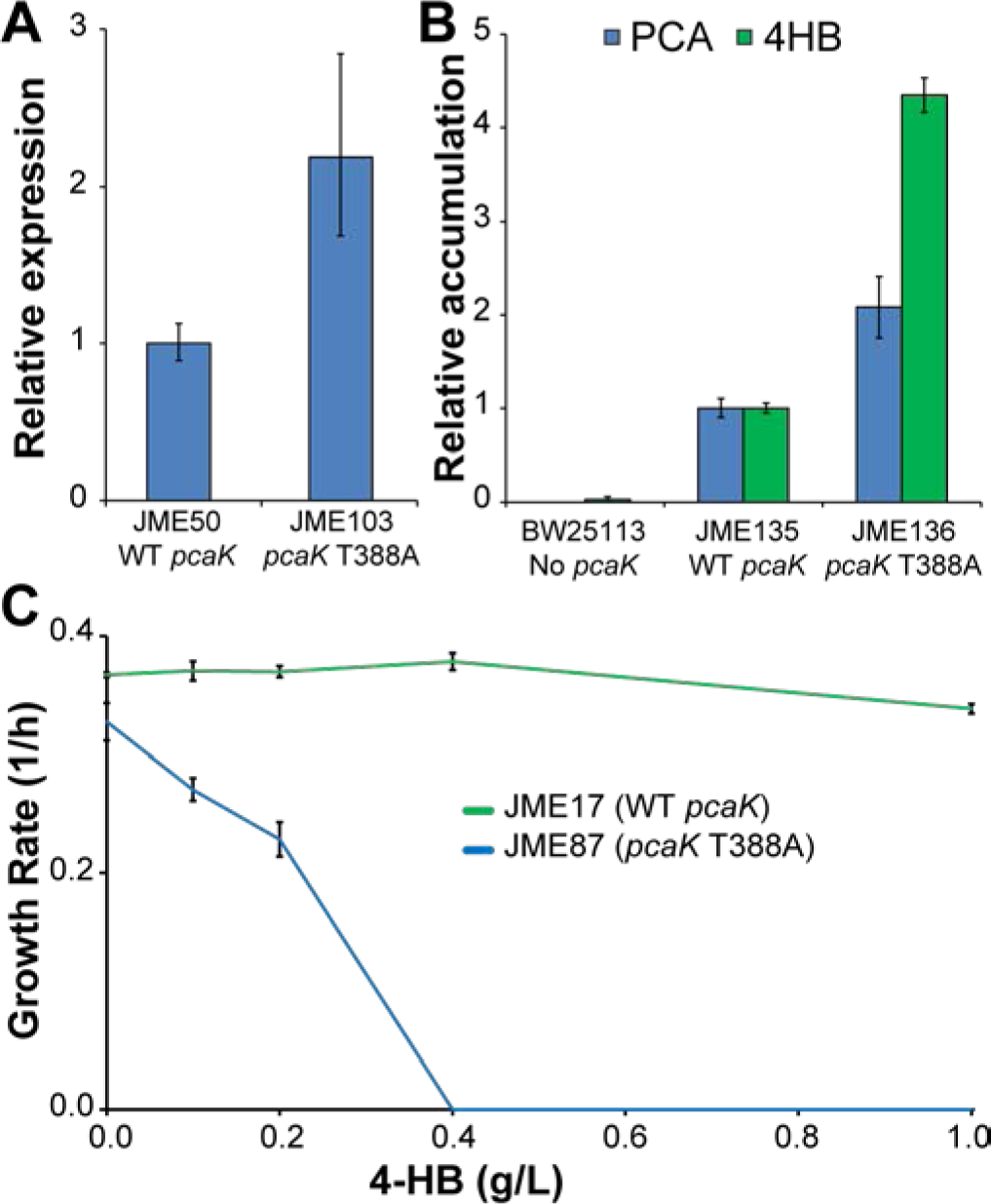
A mutation to *pcaK* affects transport of 4-HB. (A) PcaK abundance was measured in two strains that differ only in a point mutation (T388A) to *pcaK.* (B) The intracellular concentrations of PCA and 4-HB wer measured by GC/MS in strains with no transporter, the wild-type transporter, or the mutant transporter. None of the strains have a 4-HB monooxygenase or the *pcaHGBDC* operon, and therefore are unable to catabolize PCA or 4-HB. (C) Strains JME17 and JME87 were grown with PCA as the sole source of carbon and energy in media containing various concentrations of 4-HB. The two strains differ by a single point mutation to *pcaK* in JME87. Neither strain contains a 4-HB monooxygenase, preventing catabolism of 4-HB. Error bars in A-C show one standard deviation, calculated from three biological replicates. Lines in C are a guide for the eye.

We next sought to understand the effect of this mutation on transport. We first measured the intracellular concentration of PCA and 4-HB in strains that can import both compounds but cannot metabolize either. Relative concentrations were measured using GC/MS in cells exposed for 5 min to 1 g/L of PCA or 4-HB, then quickly harvested by centrifugation and washed. We found that the mutant transporter led to increased accumulation of both PCA and 4-HB, relative to the wild-type transporter, and that the increase was greater for 4-HB (Figure 5B). No significant accumulation was seen in cells lacking PcaK. PCA and 4-HB compete for transport by PcaK, so the addition of 4-HB can prevent the import of PCA (Nichols and Harwood, 1997). Using strains that can import both PCA and 4-HB but can metabolize only PCA, we measured the inhibitory effect of 4-HB (Figure 5C). In the strain with a wild-type copy of *pcaK*, the addition of 4-HB has no effect. However, 4-HB prevents growth with PCA in the *pcaK* mutant. No inhibitory effect was seen for either strain during growth with glucose. In combination, these results are consistent with the hypothesis that growth inhibition with PCA in the *pcaK* mutant was due to competition for transport rather than inherent toxicity of 4-HB (Supplementary Figure 4).

### 3.6 Functional accommodation to enzyme homologs can be homolog-specific

Two different genetic solutions can rescue growth with 4-HB in the original engineered strains, either mutations in *pcaK* or duplication of *pcaHGBDC.* We have shown that the mutations to *pcaK* increase both the expression of PcaK and its affinity for 4-HB. Conversely, duplication of *pcaHGBDC* increased expression of PcaHG and sustained rapid growth at lower concentrations of PCA. In combination, these results suggest that PcaHG has low catalytic activity in E. coli. When the internal PCA concentration is high, such as during growth with 1 g/L of PCA, the level of PcaHG activity present in JME17 is sufficient. However, at low external PCA concentrations or when the PCA is produced intracellularly by oxidation of 4-HB, the intracellular concentration will be much lower. The two genetic solutions, therefore, represent two separate biochemical solutions to increase the flux from 4-HB to PCA, either increasing transport of 4-HB or increasing expression of PcaHG (Figure 6).

Increased transport of 4-HB is sufficient for strains with either *pobA* or *praI.* Increased transport would act to raise the intracellular 4-HB concentration and these results suggest that, at a higher 4-HB concentration, both enzymes are capable of generating sufficient flux to PCA to support rapid growth. Conversely, increased expression of PcaHG lowers the threshold for flux to PCA. Increased expression of PcaHG rescues only *pobA* strains, suggesting that PraI might have a higher K_M_ for 4-HB and be unable to meet even this reduced threshold using the wild-type PcaK.

**Figure 6:**
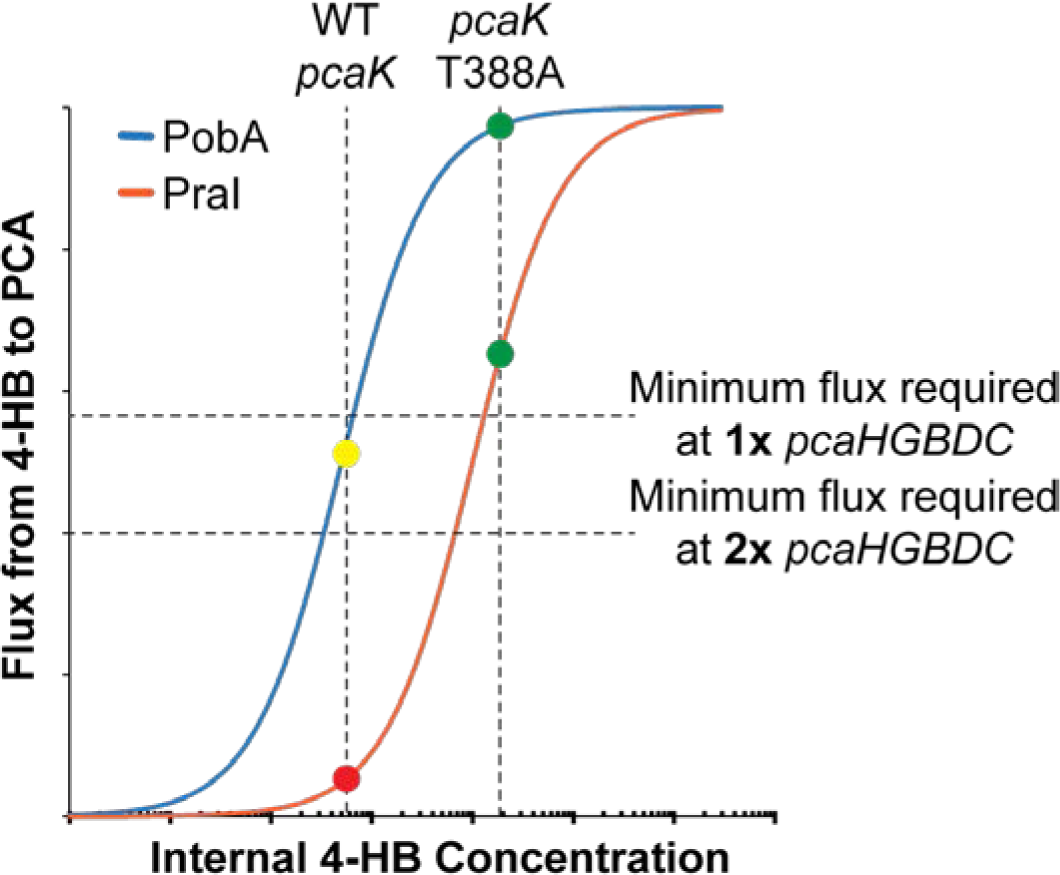
Homologous enzymes require different solutions to enable rapid growth with 4-HB. A model is shown in which mutations to the PcaK transporter increase the internal 4-HB concentration (vertical dashed lines), while modifications to the core PCA catabolism pathway allow growth at lower PCA flux (horizontal dashed lines indicate minimal flux to sustain growth). The mutant transporter thus allows growth with any combination of monooxygenase and *pcaHGBDC* copy number (green circles). The wild-type transporter cannot supply enough 4-HB to allow PraI to support growth (red circle). However, the wild-type transporter, in combination with PobA, can support growth only if the *pcaHGBDC* duplication accommodates a reduced flux to PCA (yellow circle). Semi-log Michaelis-Menten curves for PobA and PraI are drawn for illustrative purposes.

## 4.0 Conclusions

We have successfully engineered strains of *E. coli* that grow efficiently with both 4-HB and PCA, using different homologous 4-HB monooxygenases. These strains provide the foundation for effective processes to valorize lignin through initial catabolism of complex mixtures of phenolics. Our results demonstrate that standard engineering strategies can confound bioprospecting attempts by introducing shared features such as similar RNA structures. Accordingly, we showed that silent mutations to the 5’ end of the codon-optimized genes encoding 4-HB monooxygenases increased protein expression by approximately three-fold. However, increasing enzyme expression along was not sufficient to support growth. After making this change, we demonstrated that the effects on enzyme activity of mutations to the host can differ between homologs. Specifically, we identified mutations to the PcaK transporter that support growth using either the PobA or PraI enzymes, while duplication of the core PCA pathway only rescued growth with PobA. Using these methods to rapidly identify the best enzyme homolog for a given pathway, or the best genetic context for a given homolog, will greatly speed the process of metabolic pathway construction.

## 5.0 Acknowledgement

Genome resequencing and analysis was performed by Christa Pennacchio, Natasha Brown, Anna Lipzen, and Wendy Schackwitz at the Joint Genome Institute. The work conducted by the U.S. Department of Energy Joint Genome Institute, a DOE Office of Science User Facility, is supported by the Office of Science of the U.S. Department of Energy under Contract No. DE-AC02-05CH11231. Oak Ridge National Laboratory is managed by UT-Battelle, LLC, for the DOE under Contract No. DE-AC05-000R22725.

## 6.0 Funding sources

This work was supported by the BioEnergy Science Center, a U.S. Department of Energy Bioenergy Research Center supported by the Office of Biological and Environmental Research in the DOE Office of Science.

